# Characterization of zebrafish nucleus pulposus in development and aging: A fish model for probing human intervertebral disc degeneration and regeneration

**DOI:** 10.1101/2025.10.20.682618

**Authors:** Jong-Su Park, Dhivyaa Rajasundaram, Diya Patel, Zachary W. Rachfal, Kyler Hoebe-Janssen, Brian R. Cabana, Katherine A. Davoli, Kira Lathrop, Gwendolyn A. Sowa, Nam V. Vo, Xiangyun Wei

## Abstract

Intervertebral disc (IVD) degeneration (IDD) contributes to low back and neck pain. The nucleus pulposus (NP) of the IVD is usually degenerated first in age-related IDD. Addressing NP degeneration remains challenging due to insufficient understanding of its etiologies. Here, we characterize zebrafish IVD, particularly the NP, and note several characteristics that are common to or different from those of mammals. The adult zebrafish NP can be partitioned into three sections: the NP envelope with a monolayer of basal cells, the peripheral region with clustered vacuolated and non-vacuolated NP cells, and the central fibrotic region with low cellularity. With age, the percentage of vacuolated NP cells decreases, whereas that of non-vacuolated NP cells increases, as in mammals. However, a cluster of vacuolated cells persists in the NP periphery even in aged zebrafish. Notably, the ratio of the span of the fibrotic NP center to NP diameter remains largely constant at 50% regardless of age, suggesting that the zebrafish NP can better maintain its overall shape and structure, compared to mammalian NP. The fibrotic NP center and cell-populated NP periphery serve as internal controls to each other for studying NP maintenance and fibrosis. With external development, high fecundity, and great regenerative capabilities, zebrafish may be a unique NP model to complement mammalian models for studying how the NP degenerates and how the dormant capability of NP regeneration may be revitalized in humans.

## Introduction

In vertebrates, the intervertebral discs (IVDs) join the vertebrae and render the vertebral column flexible. The IVD also acts as a shock absorber to cushion physical loads imposed on the vertebral column during movements and under body weights. The functions of the IVDs are compromised in IVD degeneration (IDD) (Adams and Roughley, 2006). IDD affects two-thirds of the human population; IDD is a major cause of low back and neck pain and limits patients’ mobility (Choi et al., 2008; Peng and DePalma, 2018; Raj, 2008). However, there is no cure and few good prevention strategies for IDD (Kos et al., 2019); current treatments for IVD defects are invasive and unsatisfactory due to low efficacy and high risk for serious complications (van Uden et al., 2017). As a result, the healthcare cost of low back and neck pain is among the highest in all medical conditions (Vos et al., 2012). This status quo is difficult to change due to an insufficient understanding of the basic biology of IVD development, aging, and degeneration.

The central component of the human IVD is the nucleus pulposus (NP). The NP is gelatinous and elastic and plays a vital role in absorbing mechanical stresses imposed on the IVD. By unclear mechanisms, the NP is usually degenerated earlier in age-related IDD than other IVD components. As a result, more mechanical loads are passed to the surrounding annulus fibrosus (AF) than usual, causing AF rupture and IVD herniation. To better understand how to maintain NP and prevent or even treat IDD in humans, animal models are needed. In addition, enhanced understanding of unique characteristics in development and aging in animal models may lead to novel treatment approaches.

Here, we characterize the configuration and structure of the zebrafish IVD during development and aging, with a focus on the NP. With great regenerative capabilities, zebrafish may serve as a unique vertebrate model to provide insight into human IDD and inspiration on how IVD regeneration may be promoted in humans. Our study reveals similarities and differences between zebrafish and mammalian NPs in overall tissue architecture, the compositions of cell types, progression of degenerative fibrosis, and changes in cellularity. Compared to the human NP, the zebrafish NP better maintains its overall shape and structure during aging. These characteristics make zebrafish a unique vertebrate model to study the molecular mechanisms of NP degeneration as well as maintenance.

## Materials and methods

### Animal care and maintenance

Tübingen (TU) wildtype zebrafish were raised and maintained on a cycle of 14 hours in light and 10 hours in dark. Zebrafish embryos were raised at 28.5 °C in an incubator. Juvenile fish of 20 days postfertilization (dpf) and adult fish of 3, 5, 12, 17, and 31 months postfertilization (mpf) were used in this study. Animal care and handling were in accordance with the guidelines of the University of Pittsburgh.

### JB4 histology

Vertebral column segments of 2-4 vertebrae from the middle of the caudal spine or from the thoracic region with intact IVDs were manually isolated with forceps and needles and fixed with 4% paraformaldehyde (PFA) at room temperature for 2 hours or at 4 °C overnight. Samples were dehydrated with ethanol and embedded in JB4 resin (Polysciences, Inc.), following the manufacturer’s protocol. Embedded samples were sectioned at 2 or 5 μm in thickness. Sections were collected on slides, stained with hematoxylin and eosin (H&E) and/or Alcian blue, sealed with the Permount mounting medium, and observed under an AH2 Olympus microscope.

### Fluorescence histochemistry

Fixed segments of the vertebral column were treated with a 14% EDTA solution at room temperature for 24 hours to remove calcium, infiltrated with 40% sucrose in PBS, embedded in OCT (General Data Healthcare USA, Cincinnati, OH), and cryo-sectioned at 20 μm thickness, and collected on slides (Denville; Catalog #: M1021). AlexaFluor-488-conjugated phalloidin was used to stain F-actin (Thermo Fisher Scientific, catalog #A12379; 1:300 dilution). DAPI was used to stain nuclear DNA (Thermo Fisher Scientific; Catalog # 62248; 1:200 dilution). Immunostained sections were examined and photographed with a Fluoview FV1000 confocal microscope (Olympus).

### Whole-mount staining of isolated nucleus pulposus

The caudal NPs were isolated with forceps and needles under a stereomicroscope, fixed with 4% PFA overnight at 4 °C, stained with DAPI and AlexaFluor-488-conjugated phalloidin to visualize the cell nuclei and actin, respectively, and mounted on slides for direct examination under confocal microscopy.

### Quantification of cell numbers and fibrotic NP regions

The photographs of H&E-stained JB4 plastic sections of IVDs were used to quantify the numbers of equatorial basal NP cells, vacuolated NP cells, and interior non-vacuolated NP cells by counting cell nuclei. These cells were distinguished by their positions within the NP and possession of vacuoles or not. The fibrotic NP regions were assessed by the ratio between the diameter of the central fibrotic region, marked by elevated H&E staining, and the diameter of the entire NP. Seven sections (each section was from a different fish) were quantified for each condition. The significance of the statistical difference between any two conditions was evaluated in Excel using Student’s paired t-test with a two-tailed distribution.

### RNA-seq analysis

The zebrafish NPs were isolated manually with fine forceps and syringe injection needles under a dissection microscope from young adult fish at 3 mpf and old adult at 17 mpf. Five NPs from the middle region of the caudal vertebral column of 30 fish (150 NP in total) were isolated for each age. Isolated NPs were pooled together, washed with 1x PBS on ice three times, and homogenized in the TRIzol Reagent (Invitrogen; Catalog #: 15596026). The NP RNA was isolated according to the manufacturer’s protocol, stored at -80 °C, and shipped on dry ice to Novogene for library preparation and RNA sequencing. Differential gene expression analysis was done with the EdgeR software (Robinson et al., 2010). The |log2(FoldChange)| threshold was set at >= 1. The adjusted p-value calculation was done according to the Negative Binomial Distribution model, with an adjusted p-value threshold set at <= 0.05.

## Results

### The overall structures of zebrafish IVDs

Zebrafish contain an average of 31 vertebrae (4 Weberian, 10 precaudal, and 17 caudal vertebrae) (Bird and Mabee, 2003; Ferreri et al., 2000) (Fig. 1A is adapted from (Bird and Mabee, 2003)). Although two prior reports suggested that zebrafish IVDs contain structures that correspond to the endplate, AF, and NP of mammalian IVDs (Kague et al., 2021; Lopez-Cuevas et al., 2021), the morphological characterization of the zebrafish IVDs can be improved. We first use JB4 plastic histology to further examine the structural features of the zebrafish IVDs.

**Figure 1.**
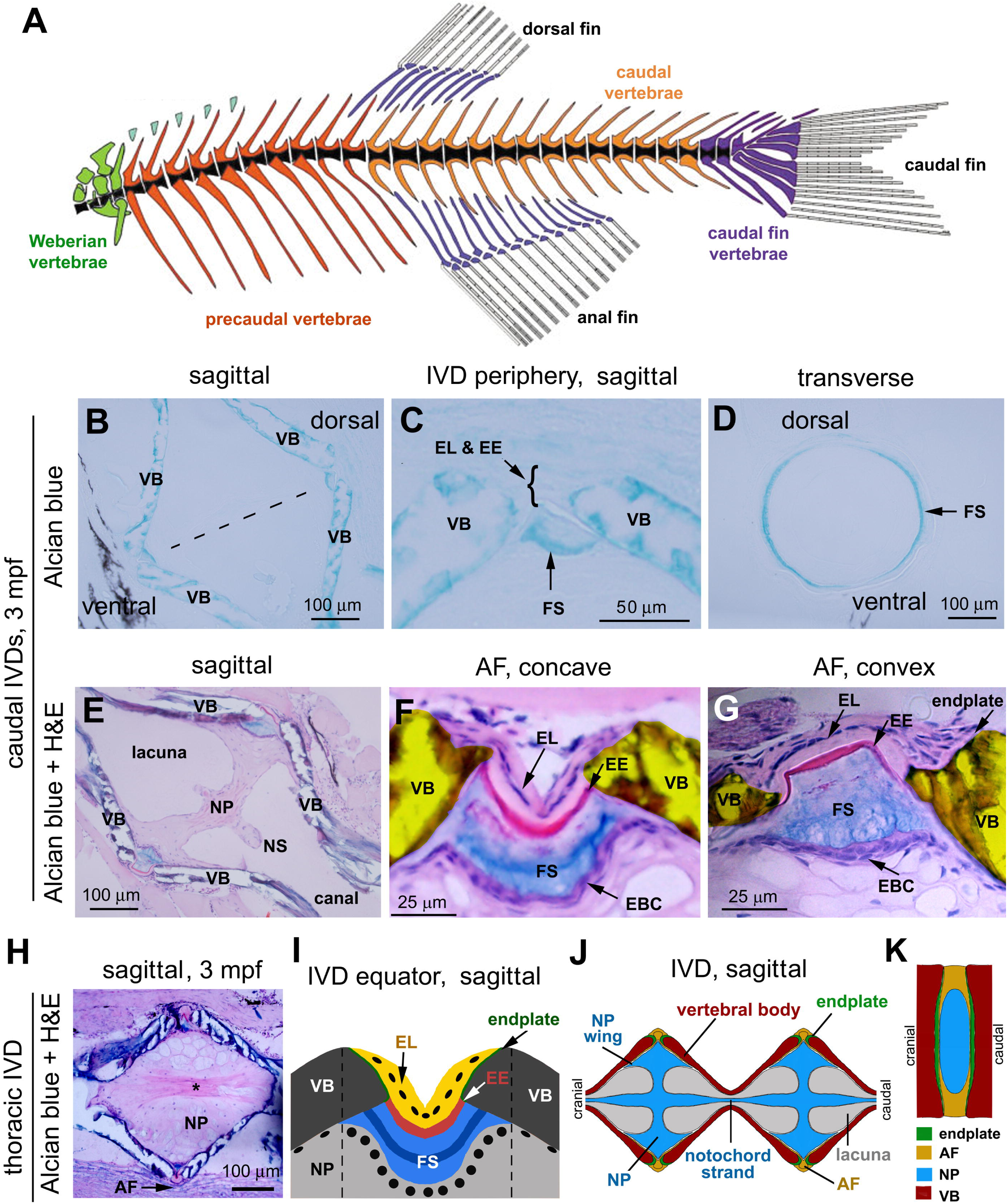
The configuration of zebrafish IVDs. **A.** A drawing of zebrafish axial skeleton, showing the division of vertebrae into four regions: Weberian, precaudal, caudal, and caudal fin. Adopted from (Bird and Mabee, 2003), with permission from the publisher. **B.** Alcian blue staining of a sagittal JB4 plastic section of a 3-mpf caudal IVD. The equatorial plane of the IVD is indicated by the dashed line. The dorsal and ventral sides of the IVD are marked. **C.** A magnification of a local region of the endplate and annulus fibrosus shown in panel B. **D.** Transverse section of the IVD at the equatorial plane, showing the circular fibrous sheath stained by Alcian Blue. **E.** Alcian Blue and H&E staining of a sagittal JB4 section of a 3-mpf caudal IVD. The intervertebral canal, lacuna, and part of the notochordal strand are indicated. **F-G.** Magnifications of the sagittal section of the IVDs, showing the endplate and AF, with the AF either in a concave (F) or convex (G) configuration. The yellow shade highlights the vertebral bodies. The endplate is sandwiched between the external intervertebral ligament and the vertebral body as a thin layer of tissue that covers the rim of the vertebral bodies. **H**. Alcian Blue and H&E staining of a sagittal section of a thoracic IVD at 3 mpf, showing that the IVD space is avoided of a lacuna and occupied entirely by the nucleus pulposus. The asterisk shows the fibrotic region in the NP center. **I.** A drawing illustrates the endplate, AF, and NP at the equatorial periphery of the IVD. Black ovals represent the cell nuclei in the external intervertebral ligament layer or the basal NP cells. Two dashed lines mark the boundaries of the equatorial region of the IVD. **J**. A diagram illustrates the sagittal view of the zebrafish caudal IVD components and the connection between two adjacent vertebrae. **K.** A diagram to illustrate the structure and configuration of mammalian IVDs. AF, annulus fibrosus; EBC, equatorial basal cells; EE, elastic externa; EL, external intervertebral ligament; FS, fibrous sheath; NP, nucleus pulposus; NS, notochordal strand; VB, vertebral body.

The surfaces of both the cranial and caudal ends of zebrafish vertebral bodies are concave (Fig. 1B, E, J). As a result, the adjacent vertebral bodies juxtapose at their rims and enclose a biconical (or amphicoelus) intervertebral space (Fig. 1C, F, J). The zebrafish endplates localize next to the rims of the cranial and caudal ends of the vertebral bodies, thus taking on a ring shape. The endplate is very thin and different from the underlying bony vertebral body, suggesting the difference in their compositions (Fig. 1F, G); a similar difference was observable in *Perca flavescens* (Schmitz, 1995). Due to their thinness, the endplates are not always easily recognizable or may be overlooked in histology. Although the ring-shaped fish endplates differ structurally from the plate-like endplates in mammals, they serve the same function of connecting the adjacent vertebral bodies through connective tissues.

Three concentric layers of connective tissues join the opposing zebrafish vertebral bodies at the rim. The outermost layer is cellular, with their cell nuclei positioned in the middle of the layer, as revealed by the blue-purple basophilic nuclear staining with hematoxylin and eosin (H&E) (Fig. 1F, G). This outermost connective tissue, which covers most of the endplate surface (Fig. 1F, G), corresponds to the external intervertebral ligament as referred to in other fish species (Schmitz, 1995; Symmons, 1979). Interior to the external intervertebral ligament, a thin layer of eosinophilic acellular structure stains distinctively in bright pink with H&E, and its strongly stained portion extends between the inner edges of the two connected endplates and tapers off drastically in the opposite direction near the inner surface of the vertebral bodies (Fig. 1F, G).

Judging by its position, this pink structure should correspond to the elastic externa observed in other fish species (Schmitz, 1995; Symmons, 1979). Further interior to the elastic externa is an Alcian Blue-philic connective tissue, which usually displays stronger staining in its central lamina than the flanking outer and inner regions (Fig. 1C, D, F, G). This Alcian Blue-philic layer is also acellular and should correspond to the fibrous sheath layer observed in other fish (Schmitz, 1995; Symmons, 1979). Together, the elastic externa and the fibrous sheath layers cover a narrow region of the inner surface of vertebral bodies adjacent to the rim in zebrafish (Fig. 1F, G), whereas they extend to cover more inner surface of the vertebral bodies in *Perca flavescens* (Schmitz, 1995). These three layers of connective tissues are very flexible; they can be stretched, compressed, either in a concave or convex configuration, depending on the nature of mechanical stresses (Fig. 1E, F, and G). Collectively, these connective tissues localize at the equatorial region of the IVD and constrain the NP to the center of the IVD as the mammalian AF does (Fig. 1I-K). Thus, they have been referred collectively to as the zebrafish AF (Kague et al., 2021; Lopez-Cuevas et al., 2021).

Interior to the AF, a septum separates the adjacent vertebrae (Fig. 1E, J; caudal spine region). This septum is the zebrafish NP, which is derived from the notochord (Haga et al., 2009; Kague et al., 2021; Lopez-Cuevas et al., 2021). This notochord–NP transition trajectory echoes that of mice (Choi et al., 2008; McCann et al., 2012) and is consistent with the fact that some NP cells contain vacuoles, a characteristic of the notochordal cells as well. At the peripheral regions of the NP, the tissue extends cranially and caudally to underlie more interior surfaces of vertebral bodies. We name these extensions the NP wings (Fig. 1E, J). Unlike being isolated in mammals, the NP of adjacent IVDs of zebrafish, or teleost fish in general, are connected at the center on both the cranial and caudal ends via the notochord strand, which is also derived from the notochord (Fig. 1E, J). Each notochord strand passes the canal at the center of a vertebra (Fit. 1E, J) (Haga et al., 2009). Therefore, the notochord strands alternate with the NPs along the craniocaudal axis of the vertebra; they connect as one single entity along the entire vertebral column. In each vertebra, the notochord strand is usually surrounded by a liquid-filled extracellular lacuna (Schmitz, 1995; Symmons, 1979) (Fig. 1E, J). However, not all vertebrae contain a lacuna; the volumes of the lacunae vary among vertebrae along the craniocaudal axis, with precaudal vertebrae containing smaller or no lacunae (Fig. 1H, a thoracic IVD) and the caudal vertebrae containing large lacunae (Fig. 1 E, J). The reduced lacuna space in the thoracic vertebrae is occupied by expanded NP as suggested by the histology (Fig. 1H). As a result, the IVD space maintains the biconical shape throughout the vertebral column, either with the presence of the lacunae or not.

The liquid-filled extracellular lacunae do not emerge from the beginning during development because at 20-dpf (days postfertilization), no lacuna is confined in larval vertebrae (Fig. 2A). The liquid-filled extracellular lacuna of the zebrafish IVD may have the physical properties similar to that of the watery extracellular matrix of the mammalian NP in handling hydrostatic pressures, except that it is not intermingled with NP cells as in the mammals. Together, the NPs, lacunae, and notochord strands are subjected to the mechanical stresses generated in swimming when the fish body bends laterally. And they potentially pass the stresses in the craniocaudal direction to the entire vertebral column because the NPs and notochord strands are physically connected (Nowroozi and Brainerd, 2012; Schmitz, 1995; Symmons, 1979). Thus, cells in the zebrafish IVDs should also bear mechanical loads as mammalian IVD cells do.

**Figure 2.**
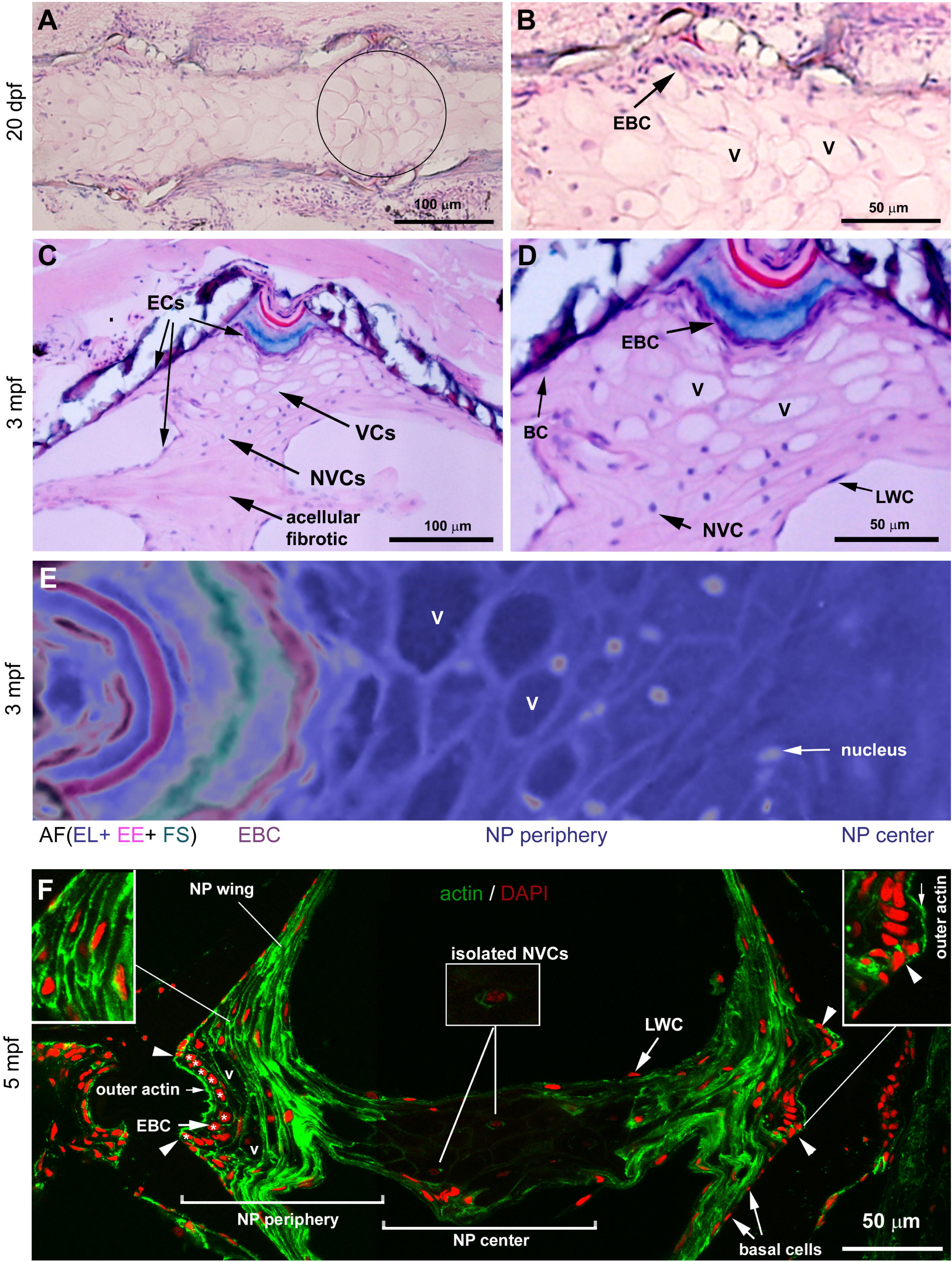
Three NP cell types in the zebrafish. **A**. Alcian Blue and H&E staining of a sagittal section of 20-dpf zebrafish vertebrae, showing that the interior of the vertebrae is filled with vacuolated NP cells (VCs). The circle arbitrarily defines the IVD region used in NP cell quantification. **B**. A higher magnification of panel A, showing the vacuolated NP cells and the equatorial basal cells (EBCs). **C**. Alcian Blue and H&E staining of a sagittal section of 3-mpf caudal vertebrae, showing the cellular NP periphery, and the fibrotic NP center. The envelope cells (ECs), vacuolated cells (VCs), and non-vacuolated cells (NVCs) are indicated. **D.** A higher magnification of panel C, showing the equatorial basal cells (EBCs), basal cells (BC), vacuoles (V) in VCs, NVCs, and lacuna wall cells (LWC). **E.** An image of a sagittal IVD section stained with Alcian Blue and H&E was rendered in Photoshop to change the colors and hues, better illustrating the regional structures and characteristics from the periphery to the center of the IVD. **F.** The sagittal images of the 5-mpf NP stained with phalloidin and DAPI to visualize actin (green) and the cell nuclei (red). The left inset shows a magnified NP peripheral region, illustrating that the local NP cells are elongated along the anterior-posterior axis of the spine. The right inset shows a transition region (arrowhead) between the EBCs and the flanking NP basal cells. The strong outer actin staining in the EBCs is indicated by an arrow. This outer actin enrichment in the EBCs abruptly reduces at the junction where the EBCs transition to NP basal cells (arrowheads). The middle inset illustrates an isolated NVC. EBCs on the left side are indicated with an arrow and asterisks.

Taken together, although the fish-specific extracellular lacunae, notochordal strands, ring-shaped endplates, and biconical-shaped IVD space (Fig. 1I) make the zebrafish IVDs look different from mammalian IVDs at first glance, the correspondence in their structural composition and functionality can be extrapolated. In particular, the zebrafish NP shows more similarities to mammalian NP, as suggested by the following analyses at the cellular and molecular levels.

### Classifications of zebrafish NP cells

To better study the NP cells, we examined the zebrafish NP by H&E staining and confocal microscopy. By H&E histology, zebrafish NP cells differ in their morphologies and locations, and they can be categorized into three groups in three NP regions: the “envelope cells” (ECs) in the surface of the NP, the “vacuolated NP cells” (VCs) and “non-vacuolated NP cells” (NVCs) in the NP periphery, and “isolated NVCs” in the central fibrotic NP region, each of which is defined and characterized as follows.

The outer boundary surface of the NP is covered by a monolayer of cells that envelops continuously the entire NP and the notochord strand. We, thus, refer to this cellular monolayer as the “NP envelope” and its constituent cells as “envelope cells” (Fig. 2C). Most of these ECs are flat; they separate the interior mass of the NP and notochord strands from the AF, the inner surface of the vertebral body, and the lacunae. These cells correspond collectively to the “basal cells” and the “lacuna wall cells” reported in *Perca flavescens* NP (Schmitz, 1995), with the former facing the AF and vertebral body and the latter facing the lacuna. In the equatorial surface of the NP, interior to the AF, the basal cells exist at higher densities than the flanking flat basal cells in the NP wings (Fig. 2A-D; Fig. 1F, G); around 20 such basal cells can be identified in each sagittal section, occupying largely the same width of the fibrous sheath layers of the AF in the craniocaudal direction. Given their equatorial position, we name these ECs “equatorial basal cells” (EBCs). Due to the higher density, EBCs are more cubical and their cell nuclei are often round; by contrast, the flat squamous basal cells in the NP wings possess flattened cell nuclei (Fig. 2D). EBCs display stronger cytoplasmic H&E staining than the flanking basal cells and interior NP cells, suggesting that they possess a distinct molecular composition. The difference in basal cell densities across the NP surface emerges even at the larval stage of 20 dpf before the lacunae emerge (Fig. 2B) and become more distinguishable at 3 mpf (Fig. 2C, D). These morphological features revealed by H&E histology suggest that the ECs, particularly EBCs, are distinct from other interior NP cells (Fig. 2D).

The interior of the NP can be divided into two regions: the “NP periphery” and the “NP center”. The periphery takes the ring shape and occupies the outer half of the NP radius, when viewed in transverse section across the AF. The NP periphery also encompass the interior of the NP wings. The remaining central NP is named the NP center, which occupy the inner half of the NP radius. In the NP periphery, exist two types of NP cells, i.e., the VCs and NVCs. VCs possess intracellular vacuoles, whereas NVC do not. VCs cluster at the outer part of the NP periphery, immediately adjacent to the EBCs and often extend into the NP wings (Fig. 2A-E). NVCs cluster at the inner part of the NP periphery, and some of them can be found at the distal end of the NP wings and within the notochord strand (Fig. 2D, E). The lack of intracellular vacuoles makes NVCs resemble mammalian chondrocyte-like NP cells. In the NP center, sparse isolated NVCs are also present.

Since NP cells are exposed to mechanical stress during swimming due to the tail bending, the stress fibers are expected to be prominent in the NP cells to cope with mechanical stress. Stress fibers are enriched with actin and myosin. We, therefore, visualized actin distribution in the NP with phalloidin under confocal microscopy. Such actin staining also highlights cell shapes and orientations (Fig. 2F). We found that actin is unevenly distributed within the NP. The strongest actin expression occurs in the NP periphery; cells in these regions are highly clustered and are elongated mostly along the anterior-posterior axis of the spine, suggesting that the mechanical stresses may be distributed along this direction (left inset in Fig. 2F). However, peripheral NVCs adjacent to the NP center are elongated radially along the NP radius. In contrast to the strong actin staining in the NP periphery, NVCs in the NP center show the weakest or no actin expression; these cells are often isolated from each other, thus referred to as the “isolated NVCs” (one of such cells is magnified in the middle inset, Fig. 2F).

ECs also display actin staining, consistent with their expected structural role in enclosing NP cells as one entity. In EBCs, actin is highly enriched at the outer side of these cells, presumably forming a cortical layer that underlies the AF-facing cell membranes. This outer actin enrichment is abruptly reduced beyond the transition point between EBCs and the flanking basal cells (Fig. 3F; the right inset; arrowheads), suggesting again that EBCs are different from other NP basal cells.

**Figure 3.**
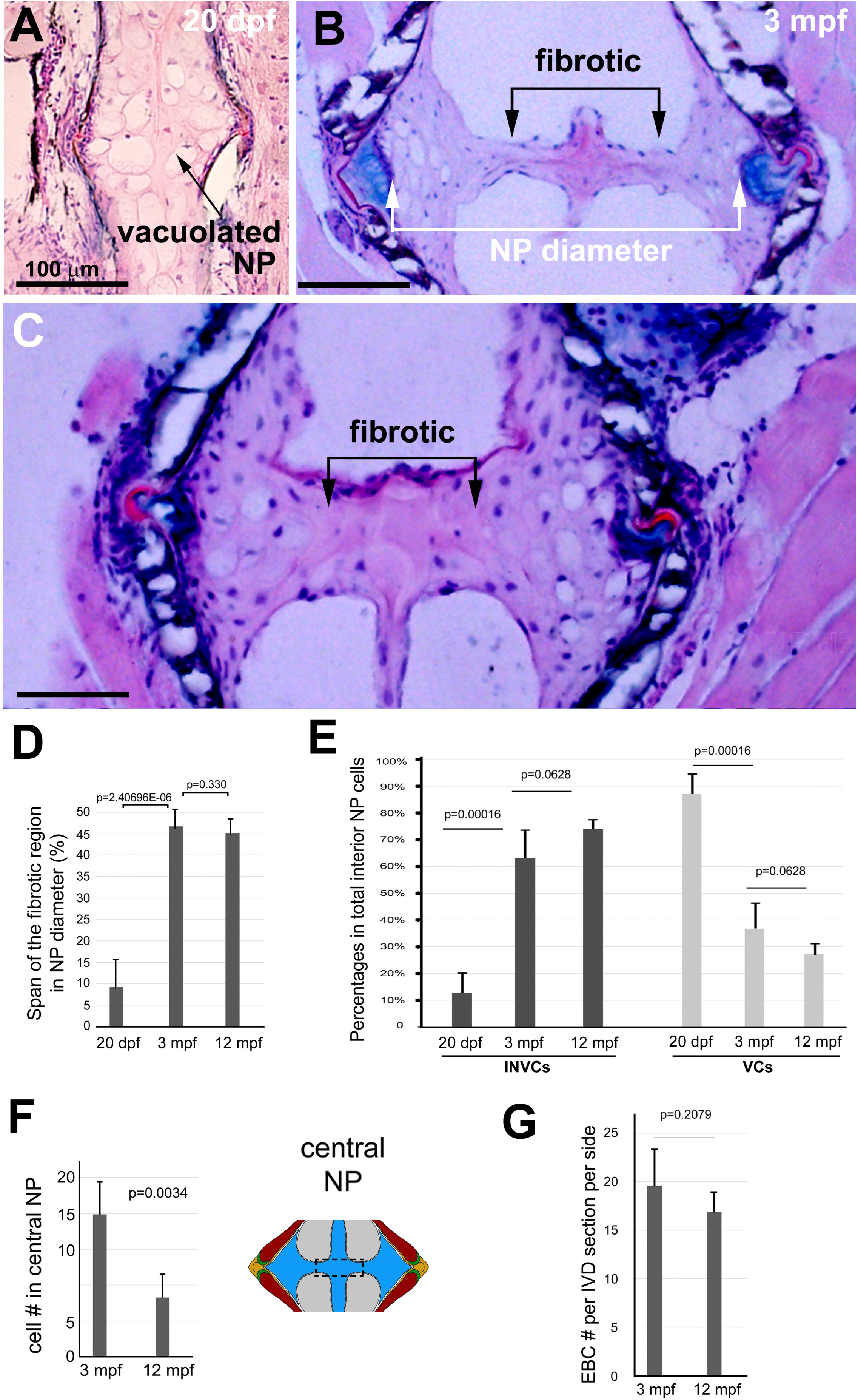
H&E staining reveals fibrotic materials in the zebrafish NP. **A.** At 20 dpf, no apparent fibrotic materials accumulated in caudal IVDs. **B, C**. Prominent fibrotic materials were detectable in the center of the NP at 3 mpf (B) and 12 mpf (C). **D.** The percentages of the span of the central fibrotic region in NP diameter at 3 mpf and 12 mpf. **E.** Average numbers of equatorial basal cells in an IVD section per side show no difference between 3 mpf and 12 mpf. **F**. The percentage of interior vacuolated and non-vacuolated NP cells in total interior NP cells at 20 dpf, 3 mpf, and 12 mpf shows age-related changes. **G**. The average number of interior NP cells in the central third region of the NP (boxed region in the IVD drawing). Seven IVD sections for each age group were analyzed.

In sum, three types of cells exist in adult zebrafish NP. Along the radius from the NP center to the equatorial surface, these cells segregate into three zones: isolated NVCs in the NP center, clustered NVCs and NCs in the periphery, and ECs in the NP envelope with EBCs as a subclass of ECs at the equator region (Fig. 2E; Fig. 2F).

### Fibrosis and changes in cellularity in the zebrafish NP with age

Abnormal accumulation of the extracellular matrix and reduction in cellularity are hallmarks of fibrotic degeneration in many tissues, such as the liver, lung, and kidney (Lopez-Otin et al., 2023; Ricard-Blum et al., 2018; Selman and Pardo, 2021). NP fibrosis is also prominent in human IDD (Peng et al., 2009; Sun et al., 2022). Thus, the degree of fibrosis is an important parameter to assess the functional status of tissues. We next examined the NP for regions that are both of low cellularity and stronger H&E staining of the extracellular matrix, manifesting as fibrotic materials. Such regions are likely regions of fibrosis. We found that the scope of such a region progresses during development (Fig. 3 A-D). At the larval stage at 20 dpf when the interior of the NP is filled with VCs, H&E staining showed little presence of fibrotic materials (Fig. 3A). However, in young adults at 3 mpf, the center of the NP displays prominent fibrotic structures stained by H&E (Fig. 3B). These stained structures are often associated with isolated NVCs, suggesting that the ECM is intermingled with these NP cells (Fig. 3B). At the 12 mpf middle-aged adult stage, the NP center is still filled with H&E-stained fibrotic structures (Fig. 3C). The ratios between the span of the fibrotic region across the NP diameter expand rapidly during development from 20 dpf to 3 mpf, but this ratio of fibrotic region to the entire NP remained roughly unchanged between 3 mpf and 12 mpf (Fig. 3D). This fibrotic appearance in the NP center was also observed in many other teleost fish (Schmitz, 1995; Symmons, 1979), suggesting that fibrosis in the NP center is common among teleost fish.

With age, the percentages of VCs among total interior NP cells reduce from over 80% at 20 dpf to below 30% at 12 mpf (Fig. 3E). Despite this reduction, some VCs persist in the periphery of the NP. Conversely, the percentages of NVCs among all interior NP cells increased from less than 20% at 20 dpf to over 70% at 12 mpf (Fig. 3E). In particular, at the adult NP center where fibrotic materials are enriched, the number of NVCs reduced significantly with age (Fig. 3F). In contrast to the drastic changes in rates of VCs and NVCs, the number of EBCs per section remains largely unchanged between 3 mpf and 12 mpf (Fig. 3G). Thus, the degrees of local fibrosis in the zebrafish NP are inversely correlated to NP cellularity.

### Changes in collagen gene expression with age

Fibrosis is normally marked by excessive deposition of collagen fibrils, and the types of collagen can also change (Intengan and Schiffrin, 2001; Ricard-Blum et al., 2018; Wynn and Ramalingam, 2012). Collagen expression changes prominently during mammalian NP degeneration. Thus, the amount of collagen deposition and types of collagen genes expressed are important parameters to assess the fibrotic state of the zebrafish NP. To analyze the changes in collagen gene expression of the NP, we performed bulk RNA-seq technology to assess the transcriptomes of NPs. This approach is doable because zebrafish NP can be isolated easily and cleanly with forceps and needles (Fig. 4A). The isolated NPs have a clear boundary, presumably due to the strength of the NP envelope. The integrity of the isolated NPs was verified by confocal imaging of actin and DAPI whole-mount staining (Fig. 4B, D), which showed NP regions that matched explicitly the corresponding regions revealed by actin and DAPI staining of in situ sections of IVDs (Fig. 4C, E). These two staining methods both revealed the NP envelope, periphery, and center in young and old adults. The ratio between the diameter of the actin-poor center and the NP diameter remains roughly at 50%, showing no drastic differences between ages and between males and females (Fig. 4G), although the absolute size of the actin-poor region increases with age, as the entire NP does. The central actin-poor NP region is only slightly larger than the central fibrotic region highlighted by H&E staining (Fig. 3D), suggesting that they correspond to each other. This actin-poor region is not yet present in 40-dpf juveniles; instead, the central region contains large VCs with strong actin staining (Fig. 4F). These results suggest that the central fibrotic region develops quickly from almost non-existent in juveniles to about 50% span of the NP diameter as observed at 3 mpf.

**Figure 4.**
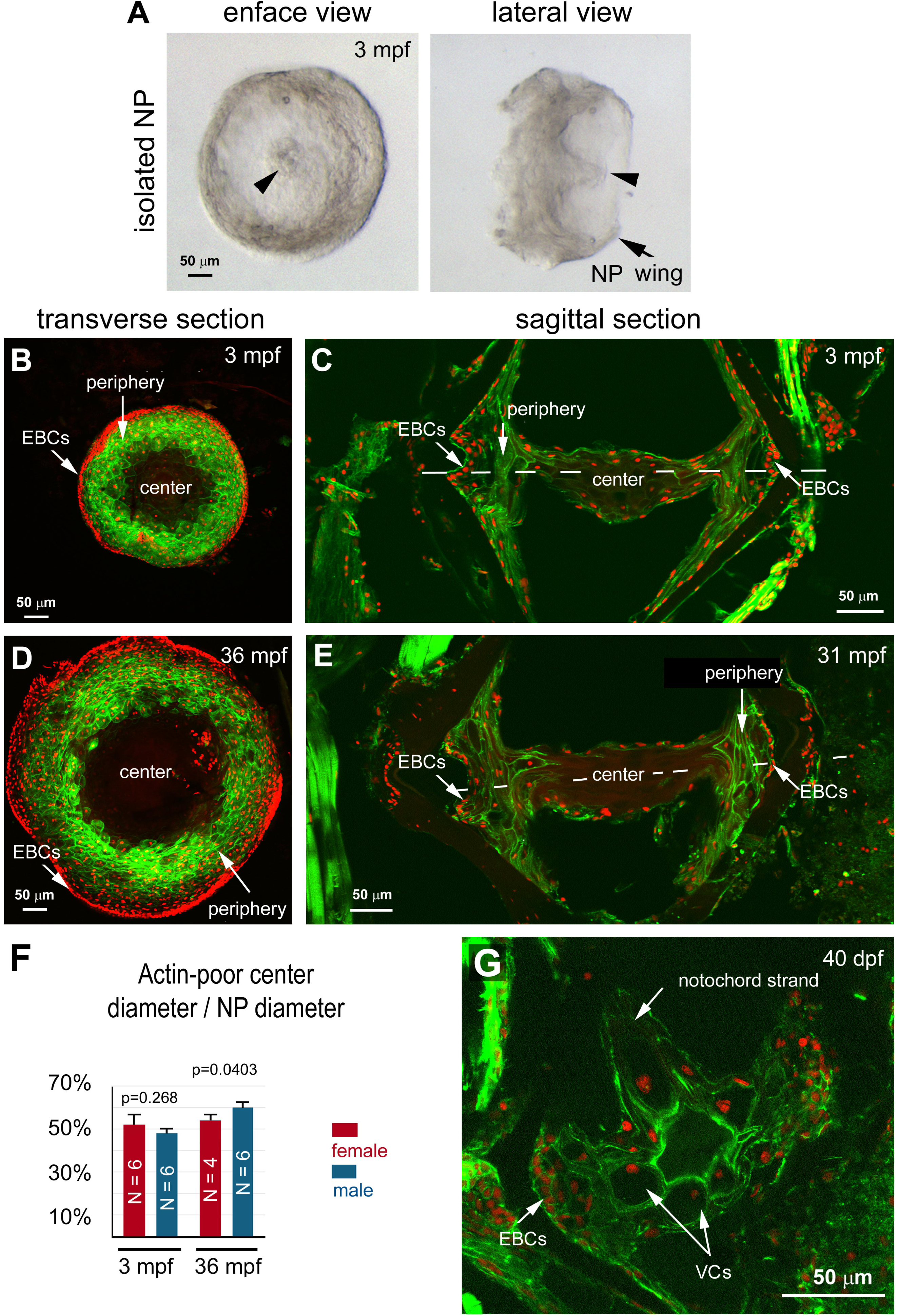
Isolation of intact zebrafish NPs. Actin and the cell nuclei were stained with phalloidin (green) and DAPI (red), respectively. **A.** Enface and lateral views of manually isolated NP under bright-field microscopy. The residues of notochord strand (arrowhead) and the NP wings are indicated. **B.** A transverse view of an isolated 3-mpf NP through the equator of the NP. **C.** A sagittal view of an IVD section at 3 mpf. **D.** A transverse view of an isolated 36-mpf NP through the equator of the NP. **E.** A sagittal view of the IVD at 31-mpf. The EBCs, peripheral region, and central fibrotic region are marked in both transverse and sagittal sections for easy comparison. **F.** The percentage of the diameter of the central actin-poor NP region in the diameter of the entire NP. The number of isolated NPs is noted in each corresponding bar. P-values between males and females of the same age were determined by Student’s t-test. **G.** An actin/DAPI staining of a sagittal section of an NP of a 40-dpf juvenile, showing the lack of an actin-poor region. VCs, vacuolated cells.

The bulk RNA-seq analysis of 3-mpf and 17-mpf NPs revealed that over 23,000 genes or variants were expressed. Among these genes, 1,588 genes were up-regulated in 17-mpf NPs, and 1,709 genes were down-regulated when compared to 3-mpf NPs (Fig. 5A). Among the expressed genes, 62 collagen genes and variants are expressed, including collagen II, IIX, IX, XI, and XVII genes that are expressed with FPKM values above 100, much higher than other collagen genes (Fig. 5B; Tables 1, 2). Many collagen genes are also differentially expressed with age. For example, collagen I was upregulated, and collagen II was downregulated with age (Tables 1, 2). These changes in collagen expression in the zebrafish NP are consistent with the presence of fibrotic material revealed by the histological H&E staining, echoing the trends of collagen expression changes in NP degeneration in mammals.

**Figure 5.**
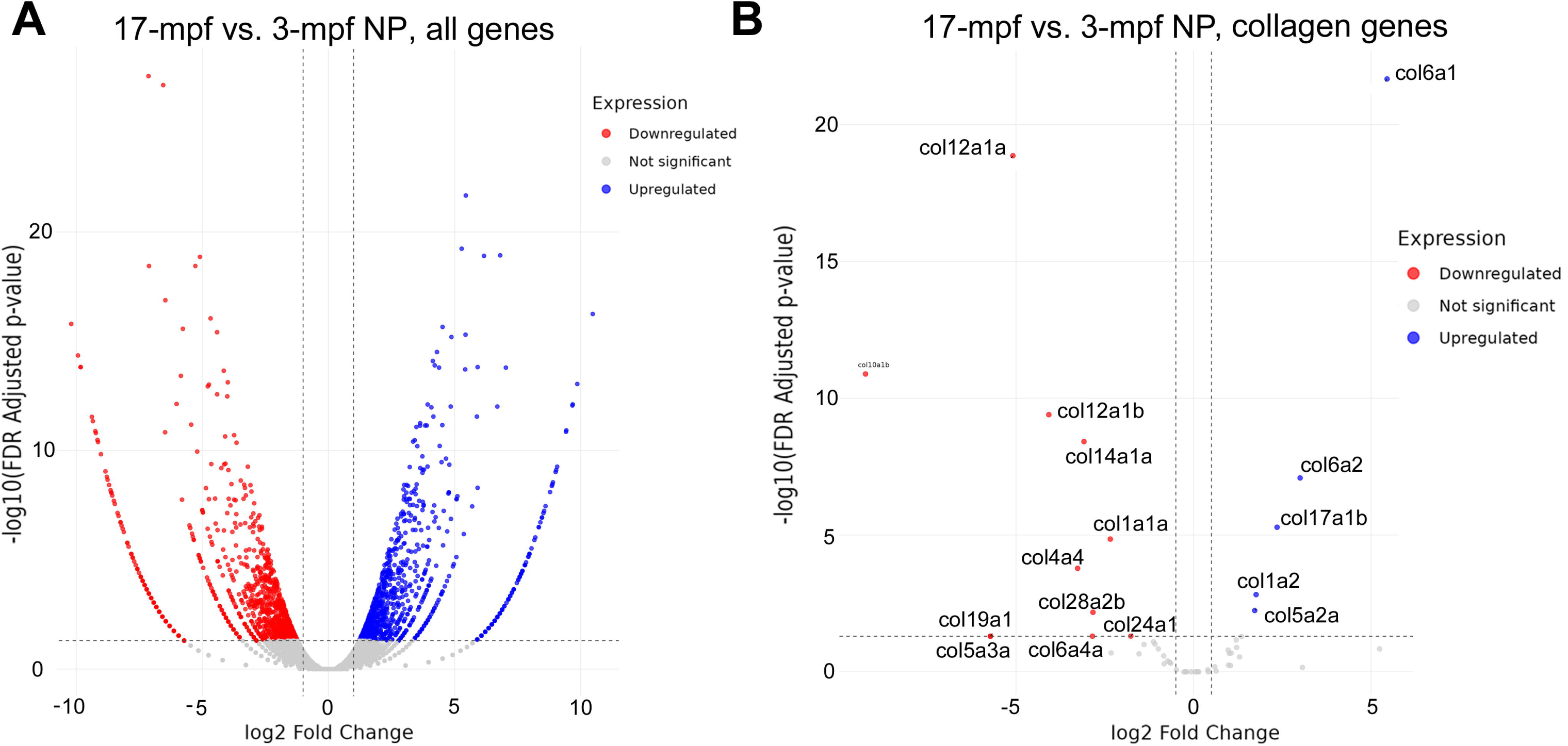
Bulk RNA-seq analysis of the isolated NPs of 3 mpf and 17 mpf. **A.** The volcano plots of the entire set of detected genes. **B.** The volcano plots of identified collagen genes.

**Table 1.** A list of NP collagen genes expressed at the level above 1 FPKM (fragments per kilobase of transcript per million fragments mapped). Rows in blue indicate NP collagen genes that are up-regulated at 17 mpf compared to 3 mpf, rows in red indicate down-regulated collagen genes, and rows in black are ones not significantly changed. Rows in bold indicate collagen genes expressed prominently at FPKMs above 10.

| genes | 17mpf_fpkm | 3mpf_fpkm | log2FoldChange | pvalue |
| --- | --- | --- | --- | --- |
| **col6a1** | **25.41411552** | **0.61986717** | **5.347880012** | **4.35E-25** |
| col4a3bpa | 3.453383128 | 0.14865736 | 4.506138125 | 9.96E-13 |
| col6a2 | 4.454893068 | 0.34078865 | 3.694831149 | 3.23E-12 |
| **col17a1b** | **180.2862383** | **34.7133593** | **2.370544334** | **4.05E-08** |
| col5a2a | 2.354595352 | 0.57852931 | 2.016232089 | 2.52E-05 |
| **col1a2** | **16.47716323** | **4.87558214** | **1.750357408** | **4.85E-05** |
| col15a1b | 1.649014828 | 0.55207565 | 1.569628708 | 0.00147 |
| col8a1a | 3.858946932 | 1.28581094 | 1.576402968 | 0.001532 |
| col12a1a | 0.218993646 | 7.66439976 | -5.130147727 | 7.31E-23 |
| col10a1b | 0 | 1.91429755 | -9.151568364 | 3.82E-12 |
| col14a1a | 0.952698519 | 7.24915184 | -2.932492079 | 1.47E-10 |
| **col11a1a** | **18.30748204** | **54.4797866** | **-1.579328849** | **0.000181** |
| col1a1a | 0.854339732 | 2.85565777 | -1.74519062 | 0.000188 |
| **col9a1b** | **232.4513284** | **539.516892** | **-1.220854294** | **0.003369** |
| **col9a3** | **103.0660556** | **226.186431** | **-1.140047493** | **0.006143** |
| col27a1a | 2.726848733 | 1.02105359 | 1.406235518 | 0.012853 |
| col4a5 | 1.008361553 | 0.43151482 | 1.215834598 | 0.015654 |
| **col8a1b** | **206.4051685** | **104.051199** | **0.982040747** | **0.017918** |
| col4a6 | 1.246224713 | 0.586097 | 1.080887649 | 0.021241 |
| **col11a2** | **71.67237198** | **133.933827** | **-0.908147266** | **0.028228** |
| col4a3bpa | 3.324413266 | 6.35532306 | -0.940409955 | 0.032399 |
| col1a1b | 3.006296432 | 5.30442998 | -0.825014631 | 0.05602 |
| col27a1b | 2.964853912 | 1.59521334 | 0.88703401 | 0.06019 |
| col15a1a | 1.012139784 | 0.44625686 | 1.167835048 | 0.069628 |
| **col2a1a** | **343.0176209** | **569.013847** | **-0.736304** | **0.074253** |
| **col9a2** | **293.6917434** | **475.097082** | **-0.700040534** | **0.089621** |
| col27a1a | 4.516822822 | 2.71706749 | 0.726629283 | 0.104986 |
| col8a2 | 3.470193545 | 2.70304157 | 0.353972673 | 0.437613 |
| **col11a1a** | **91.92831056** | **112.527631** | **-0.297820561** | **0.468834** |
| col27a1b | 2.728470173 | 3.23448428 | -0.251468514 | 0.556683 |
| col4a2 | 5.064027988 | 5.85044246 | -0.214343337 | 0.609736 |
| **col9a1b** | **181.0447034** | **163.706093** | **0.139109454** | **0.735366** |
| col4a1 | 5.073873706 | 5.30703179 | -0.070920658 | 0.873512 |
| **col4a3bpb** | **12.92679536** | **12.6488475** | **0.025228046** | **0.954511** |

**Table 2.** A list of NP collagen genes expressed at levels lower than 1 FPKM. Rows in red indicate NP collagen genes that are downregulated at 17 mpf compared to 3 mpf, and rows in black are those not significantly changed.

| genes | 17mpf_fpkm | 3mpf_fpkm | log2FoldChange | pvalue |
| --- | --- | --- | --- | --- |
| col12a1b | 0.026645937 | 0.975697165 | -5.162575406 | 3.58E-15 |
| col4a4 | 0.011064417 | 0.415879323 | -5.057956373 | 1.30E-06 |
| col28a2b | 0.045559055 | 0.478744617 | -3.341268087 | 5.75E-05 |
| col24a1 | 0.145812487 | 0.813260459 | -2.453709899 | 0.001648699 |
| col19a1 | 0 | 0.142605499 | -6.487130293 | 0.002434933 |
| col2a1b | 0.131919343 | 0.436228619 | -1.719073544 | 0.010958385 |
| col7a1 | 0.605428152 | 0.266935924 | 1.17282463 | 0.02049697 |
| col17a1a | 0.081299585 | 0.253477783 | -1.627655651 | 0.033910222 |
| col13a1 | 0 | 0.133264349 | -5.760846726 | 0.040964585 |
| col5a1 | 0.606584752 | 0.303783628 | 0.989416489 | 0.04808137 |
| col6a4a | 0.051744951 | 0.141165185 | -1.438483087 | 0.058010008 |
| col25a1 | 0.019214175 | 0.104836221 | -2.376211218 | 0.062338843 |
| col28a1b | 0.367717149 | 0.79784092 | -1.118020726 | 0.069235192 |
| col5a3a | 0 | 0.050238544 | -5.50312382 | 0.075101739 |
| col28a1a | 0.060875327 | 0 | 5.225935384 | 0.075101739 |
| col5a3b | 0.112194892 | 0.027206971 | 1.978391817 | 0.094716843 |
| col18a1b | 0.026310947 | 0.111655887 | -2.01873139 | 0.143685221 |
| col13a1 | 0.176097648 | 0.12810967 | 0.448053975 | 0.616742499 |
| col2a1a | 0.053626565 | 0.03901291 | 0.438724577 | 0.746294654 |
| col21a1 | 0.07406991 | 0.053885284 | 0.438724577 | 0.746294654 |
| col6a3 | 0.219934807 | 0.202299941 | 0.114313296 | 0.827175791 |
| col16a1 | 0.113480528 | 0.12612753 | -0.156865148 | 1 |
| col10a1a | 0.048848852 | 0.0888429 | -0.826508986 | 1 |
| col11a1b | 0.103361148 | 0.10443671 | -0.020824412 | 1 |
| col18a1a | 0.067839878 | 0.058753552 | 0.197182182 | 1 |
| col5a2b | 0.059100078 | 0.057326463 | 0.036808896 | 1 |
| col9a1a | 0.058780032 | 0.035635013 | 0.657566114 | 1 |

### Expression of stem cell markers in zebrafish NP

The observation that the ratio of the fibrotic center to the NP diameter remains roughly the same with age suggests that the zebrafish NP may be capable of maintaining a balance, at the tissue level, between NP growth (or regeneration) and fibrosis (which involves cell death as suggested by the reduction in cellularity in the NP center; Fig. 3F). Many stem cell or progenitor cell markers have been proposed to be expressed in mammalian NP, including *tnc* and *uts2r* (Gao et al., 2022), *tagln* (Tan et al., 2024), *cts12* (Chen et al., 2024), *jagged* and *notch1* (Liu et al., 2017), *nabp1a* and *mki67* (Swahn et al., 2024), tie2 and disialoganglioside 2 (Sakai et al., 2012), cell surface proteins Cd29, Cd44, Cd73, Cd90, and Cd105) (Wu et al., 2018), and transcription factors *OCT4*, *nanog*, and *sox2* (Li et al., 2020). Although the characteristics of the NP stem or progenitor cells are still to be fully understood and the reliability of these markers is yet to be agreed upon, these candidate markers provide some handles to study the molecular mechanisms of NP homeostasis. We thus examined whether the counterparts of these candidate markers are expressed in zebrafish NP. The RNA-seq analysis revealed that many markers were indeed expressed in the zebrafish NP (Table 3), although the expression *of Oct4, sox2, cd29, cd73, cd90, cd105, tie2*, and *disialoganglioside* was not detected. However, the expression levels of the candidate stem cell marker genes varied between 3 mpf and 17 mpf, and the variations do not follow a particular trend; some genes were upregulated, while others are unchanged or downregulated, with statistical significance (Table 3), suggesting a possibility of their differential involvement in zebrafish NP homeostasis.

**Table 3.**
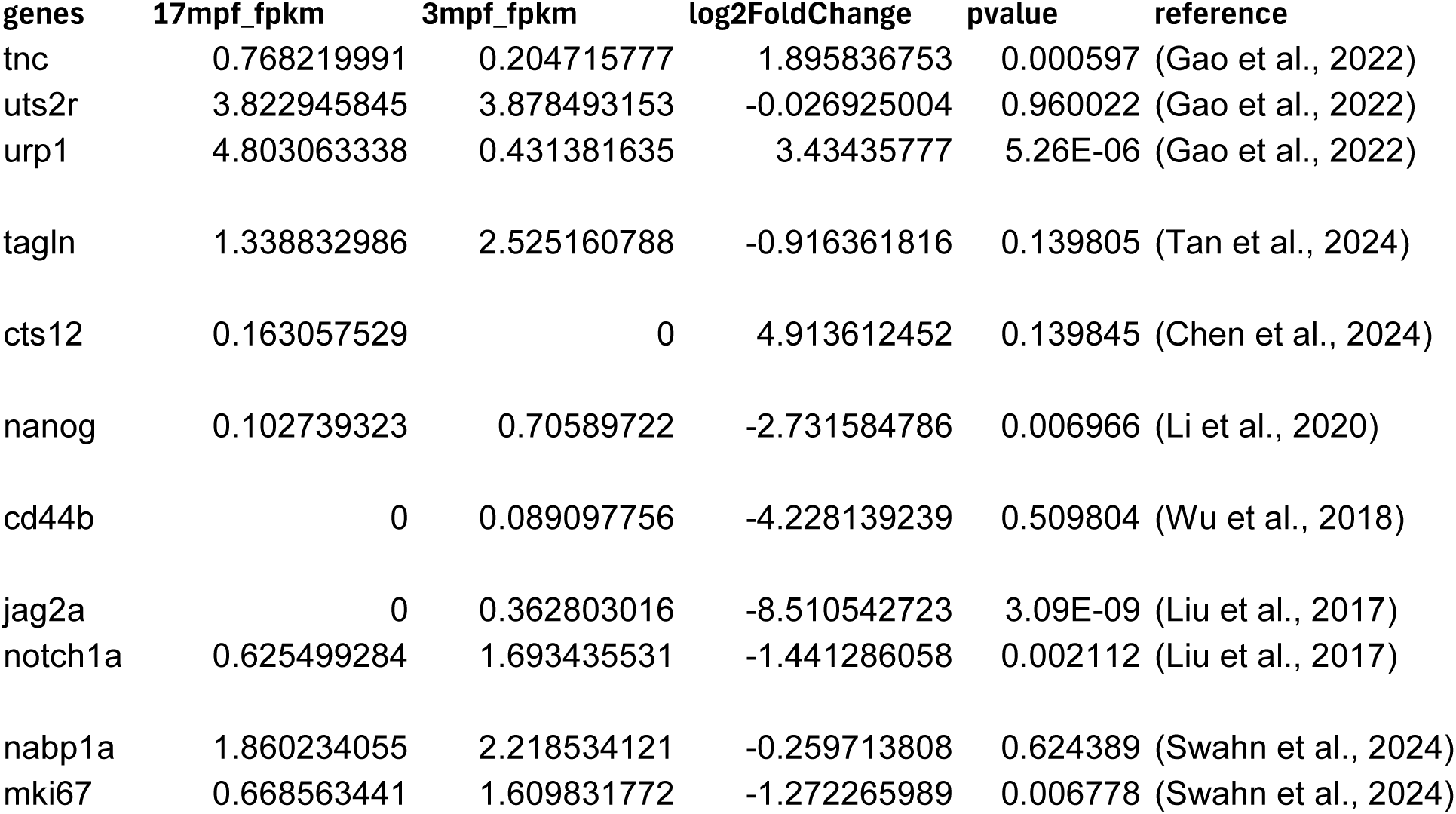
Zebrafish counterparts of mammalian NP progenitor candidate markers.

| genes | 17mpf_fpk | 3mpf_fpk | log2FoldChange | pvalue | reference |
| --- | --- | --- | --- | --- | --- |
| tnc | 0.768219991 | 0.204715777 | 1.895836753 | 0.000597 | (Gao et al., 2022) |
| uts2r | 3.822945845 | 3.878493153 | -0.026925004 | 0.960022 | (Gao et al., 2022) |
| urp1 | 4.803063338 | 0.431381635 | 3.43435777 | 5.26E-06 | (Gao et al., 2022) |
| tagln | 1.338832986 | 2.525160788 | -0.916361816 | 0.139805 | (Tan et al., 2024) |
| cts12 | 0.163057529 | 0 | 4.913612452 | 0.139845 | (Chen et al., 2024) |
| nanog | 0.102739323 | 0.70589722 | -2.731584786 | 0.006966 | (Li et al., 2020) |
| cd44b | 0 | 0.089097756 | -4.228139239 | 0.509804 | (Wu et al., 2018) |
| jag2a | 0 | 0.362803016 | -8.510542723 | 3.09E-09 | (Liu et al., 2017) |
| notch1a | 0.625499284 | 1.693435531 | -1.441286058 | 0.002112 | (Liu et al., 2017) |
| nabp1a | 1.860234055 | 2.218534121 | -0.259713808 | 0.624389 | (Swahn et al., 2024) |
| mki67 | 0.668563441 | 1.609831772 | -1.272265989 | 0.006778 | (Swahn et al., 2024) |

### Zebrafish spinal bending model to study mechanical stresses on the NP

In zebrafish, the IVDs are expected to endure mechanical stresses when fish swim in a carangiform/subcarangiform locomotion style (Vanhaesebroucke et al., 2023). This type of movement depends on the lateral bending of the caudal spine. This movement causes compressive stresses to the IVD on the concave side of the bending and tensile stresses on the convex side. To examine whether such bending induces conformational changes to the IVD, we fixed fish in bent postures as a simulation of bending occurring in swimming. We found that bending induced observable conformational changes to the NP and the AF (Fig. 6. A-C). For example, the AF was often pushed outward on the concave side of bending and sucked inward on the convex side. Furthermore, the cell nuclei of the EBCs are stretched flat on the convex side and compressed to round up on the opposite side (Fig. 6B, C). Thus, the swimming-induced mechanical stresses may similarly cause conformational changes to the IVD at both the tissue and cellular levels, although swimming-induced mechanical stress should be transient and oscillate between compression and stretching. The effects of such mechanical stresses on gene expression and IVD maintenance may be analyzed *in vivo* by positioning live fish in a body restriction mold, which subjects the spine to various degrees of curvatures to induce stresses at different levels, with higher stresses in more bent postures (Fig. 6D).

**Figure 6.**
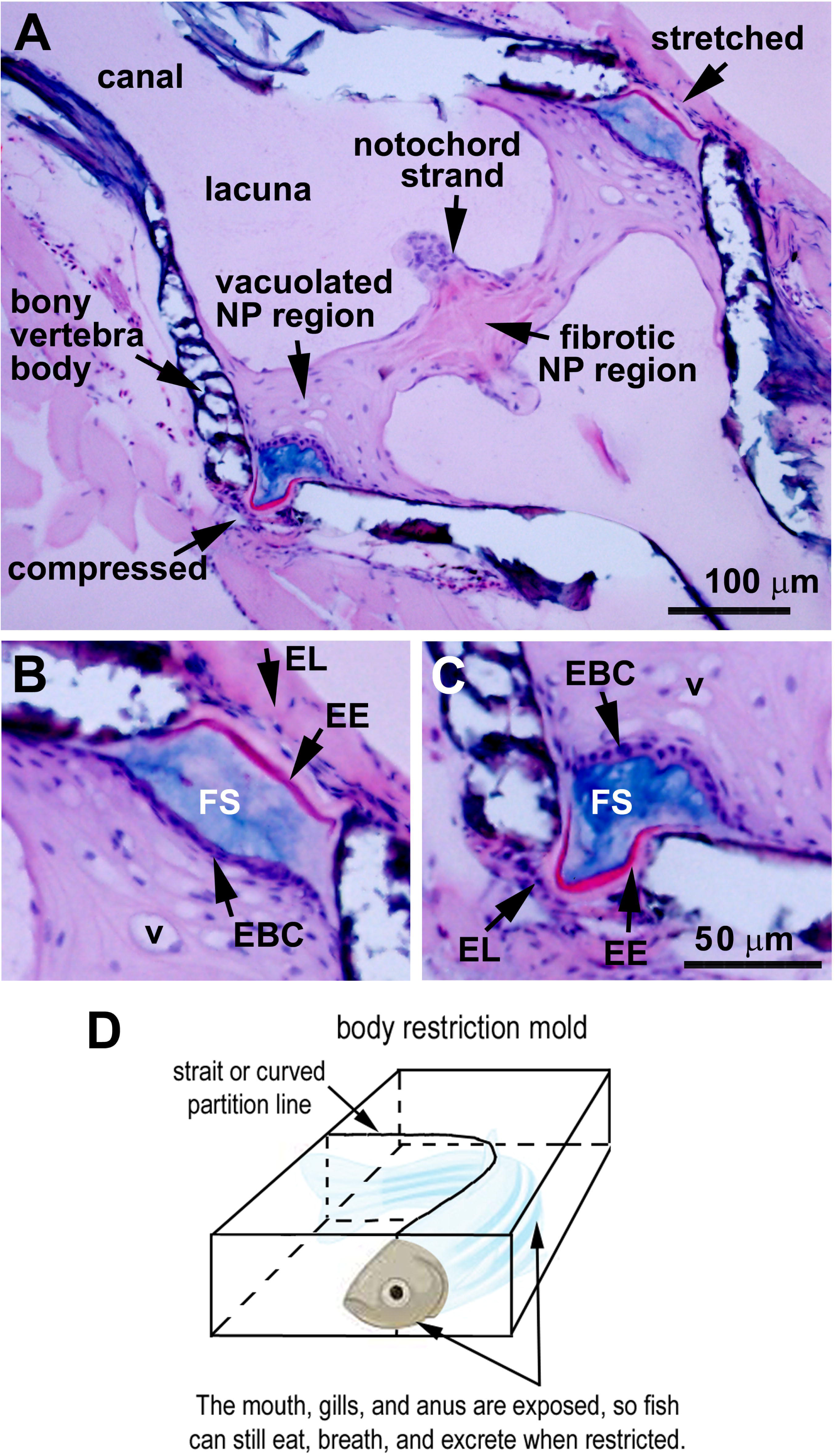
Conformational changes of the IVDs under compressive and tensile stresses. **A.** A bent 3-mpf IVD shows the AF and peripheral NP compressed on the concave side of bending and stretched on the convex side. Note that the AF was pushed outward at the compressed side and stretched flat at the convex side. **B**. A higher magnification of the stretched IVD periphery in Panel A shows the flat EBC nuclei and AF. **C.** Higher magnification of the compressed IVD periphery in panel A shows the squeezed AF layers and the round EBC nuclei. **D.** A diagram illustrates how a body restriction mold can bend fish body to cause compressive and tensile stresses to the spine in live fish.

## Discussion

The molecular mechanisms of IDD are yet to be better understood with animal models. Mammalian models have provided much knowledge, but their intrauterine development and lower fecundity make it inconvenient to study IVD development and maintenance with genetic approaches. In this study, we characterize the configuration and structure of zebrafish IVD, particularly the NP. We propose that zebrafish, with greater regenerative capabilities, are a unique model for studying NP degeneration and regeneration and may reveal novel mechanisms to be explored in humans.

### NP cell morphologies and organization in fish vs. mammals

Zebrafish and mammalian NPs are all derived from the notochord (Choi et al., 2008; Haga et al., 2009; McCann et al., 2012). The two types of mammalian NP cells, i.e., the vacuolated NP cells and the non-vacuolated NP cells, have corresponding counterparts in the interior of the zebrafish NP, and they display three similar changes in morphology and organization during development and aging. First, the percentage of vacuolated cells in total interior NP cells decreases with age, whereas the percentage of non-vacuolated cells increases drastically when fish develop into adults. Because the vacuoles may render NP cells the rigidity to directly resist physical pressures endured by the NP as notochordal cells do in the notochord, the reduction in vacuolated NP cells suggests that this capability is compromised both in fish and mammals with age. However, it is worth noting that this capability is not entirely lost in fish because a group of vacuolated NP cells persists in the outer periphery of the zebrafish NP, regardless of age. By contrast, in mammals, all vacuolated NP cells are eventually replaced with non-vacuolated cells (Hunter et al., 2003b, 2004; Sakai et al., 2009).

Second, in both zebrafish and mammals, vacuolated NP cells are clustered together, whereas non-vacuolated NP cells are more separated as dispersed individual cells, resembling chondrocytes (Hunter et al., 2003b, 2004; Sakai et al., 2009). The modes of vacuolated NP cell clustering, however, differ between fish and mammals. In mammals, vacuolated NP cells were observed to form 3D network clusters, which are embedded in the extracellular matrix. The extracellular NP matrix plays an important role in resisting compression because the matrix is composed largely of water, collagen II, and proteoglycan aggrecan (Hunter et al., 2003a), whereas in zebrafish, vacuolated NP cells form a more compact mass in the outer periphery of the NP as a ring. Nevertheless, images of many studies also showed large masses of clustered NP cells in mammals, particularly in young animals or fetuses (Sakai et al., 2012; Tan et al., 2024; Wang et al., 2021), suggesting that NP cell clustering is a universal cellular organization pattern in young IVDs. Despite the variations in the modes of NP cell clustering, the physical intercellular adhesions between clustered NP cells may be conserved at the level of cell-cell adhesion between fish and mammals. This NP cell clustering may sustain cell adhesion-based mechanosensation, which was proposed to underlie NP gene expression regulation under mechanical stresses {Wei, 2025 #5377}. This cellular clustering is lost when vacuolated NP cells are replaced with dispersed non-vacuolated NP cells. Thus, the potential cell-clustering-based mechanotransduction mechanisms are compromised in both fish and mammalian NPs during aging, with the former occurring locally in the center of the NP and the latter entirely throughout the NP.

Third, the age-related decreases in cellularity either locally in the center of zebrafish NP or entirely in mammalian NP undermine the capability of NP cells to regulate NP function in handling physical loads. In mammalian IDD, apoptosis of NP cells is one way that the NP gradually loses its cellularity (Zhu et al., 2023). Similarly, the center region of the zebrafish NP also undergoes apoptosis (Rayrikar et al., 2023), suggesting that apoptosis is also a way by which NP cellularity reduces in zebrafish during aging. As a result of this reduction in cellularity, the NP cells in the center of the zebrafish NP become more isolated, thus compromising any direct cell-cell interactions that are preserved in the periphery of the NP. In zebrafish, NP cell apoptosis can be either reduced or enhanced by overexpressing or suppressing cellular communication network factor 2a (ccn2a), respectively (Rayrikar et al., 2023). The loss of ccn2a also causes an early onset of IDD in mice (Bedore et al., 2013). This conserved role of ccn2a in promoting NP cell survival suggests that understanding how apoptosis is triggered in the center of zebrafish NP will help with understanding NP cell apoptosis in mammals.

Thus, the similarities between zebrafish and mammals in NP cell morphologies, organization, and cell maintenance suggest that similar molecular mechanisms underlie the developmental and aging trajectories of zebrafish and mammalian NP cells. On the other hand, the unique persistence of NP cell clustering in the periphery of the zebrafish NP contrasts with the eventual loss of cell clustering in mammals (Hunter et al., 2003b, 2004; Sakai et al., 2009). It would be important to determine whether this persistent NP cell clustering contributes to the prevention of fibrosis in the periphery of the NP and thus the overall NP maintenance in zebrafish.

### Fibrosis of the NP in zebrafish vs. mammals

Fibrosis is prominent in human IDD (Sun et al., 2022). This fibrotic change can drastically compromise the capability of the NP in handling physical loads because when the watery NP extracellular matrix is reduced in fibrosis, higher compressive stresses can be passed to the surrounding AF, causing its rupture and IVD herniation. Thus, preventing NP fibrosis may be one important way to halt the deterioration of the NP functions in IDD. To date, there is no good way to prevent or slow down NP fibrosis in humans.

To better understand the molecular mechanisms of NP fibrosis in humans, the center of zebrafish NP may serve as a model. We and others show that this fibrosis starts as soon as 3 mpf (Haga et al., 2009; Hayes et al., 2013). This quick development of fibrosis makes zebrafish a practical model to study age-related NP fibrosis, even though zebrafish can live 2-3 years and their reproductive capability does not start to decline until around 12 mpf. As collagen deposition contributes to fibrosis, comparing the changes in collagen expression in NP fibrosis between mammals and zebrafish would provide useful information to understand the molecular mechanisms that underlie NP fibrosis. Healthy mammalian NPs express predominantly collagen II, which makes an extracellular matrix meshwork together with other matrix proteins like aggrecan, rendering the NP gelatinous to withstand hydrostatic pressures (Eyre and Muir, 1974, 1977). In contrast, in degenerated human NP, collagen I is upregulated (Eyre and Muir, 1977; Lv et al., 2016; Song et al., 2022; Yee et al., 2016; Zeldin et al., 2020), whereas collagen II and aggrecan are downregulated (Sztrolovics et al., 1997; Yee et al., 2016). Excessive collagen I in the human NP matrix makes the NP more fibrotic, losing its gelatinous state for the shock-absorbing function (Sun et al., 2022). Similar fibrotic changes were also observed in other mammals, including mice (Au et al., 2020; Lv et al., 2016; Tessier et al., 2020; Zhang et al., 2018), horses (Bergmann et al., 2022), rabbits (Leung et al., 2014), and rats (Feng et al., 2017; Kong et al., 2021; Wang et al., 2019). Although these trends of collagen expression changes during fibrosis are also similarly observed in zebrafish (Yee et al., 2016; Zeldin et al., 2020; Zhang et al., 2018), our RNA-seq analysis suggests that it would be too simple to merely say that changes in collagen I and II are the major collagen factors contributing to the fibrotic progression in age-related NP degeneration, because, besides collagen I and II, over 60 collagen genes and variants are expressed in zebrafish NP. Supporting this notion, RNA-seq analysis of human NP also revealed the expression of many collagen gene types, including Col2a1, Col9a3, and Col11a1 (Fernandes et al., 2020). Thus, further investigation of collagen gene expression regulation in the zebrafish NP should shed light on the molecular mechanisms of NP fibrosis in humans.

### Zebrafish as a regeneration model to study NP homeostasis

Paradoxically, zebrafish NP may also serve as a regeneration model besides an NP degeneration model. Like other highly regenerative species, such as hydra, planaria, and newts, zebrafish have been regarded as the best available regenerative vertebrate model to study the regeneration of many organs, including the retina, heart, liver, etc. because zebrafish can provide insight into the dormant regeneration capabilities in humans and how to revitalize such capabilities to prevent and treat degeneration (Gemberling et al., 2013; Massoz et al., 2021; Mehta and Singh, 2019; Ribitsch et al., 2020). Zebrafish may also be a good model for NP regeneration per the following reasoning.

Evidence suggests that EBCs may harbor NP stem or progenitor cells. The basal cells in zebrafish NP may be derived from notochordal sheath cells, which surround the interior notochordal cells. This speculated origin of the basal cells is supported by two facts: First, NP basal cells and notochordal sheath cells localize at the surface to encapsulate the corresponding interior cells in either tissue. Second, they are both non-vacuolated. Highly proliferative cells were also identified in the peripheral region of zebrafish IVD by EdU incorporation recently (Rayrikar et al., 2023). The persistent equatorial basal NP cells prompt us to hypothesize that these cells may act as NP stem cells to supply new vacuolated NP cells to the NP interior, which in turn trans-differentiate into non-vacuolated cells to supplement cells to NP central regions, where non-vacuolated NP cells eventually degenerate. By so doing, the zebrafish NP can maintain homeostasis at the tissue level. Analogy to this transition, in the zebrafish notochord at early larval stages, the outer notochordal sheath cells (the suspected precursors of the NP basal cells) can move interiorly and differentiate into vacuolated notochordal cells (Garcia et al., 2017).

The above hypothesis echoes the recent identification of NP progenitor cells at the periphery of the NP in mice and humans, although these NP progenitor cells are eventually lost in IDD (Gao et al., 2022; Tan et al., 2024). NP-derived stem cells have also been isolated by culturing explants of human NP tissues (Cui et al., 2024; Wu et al., 2017) and by single-cell sequencing analyses of rat IVDs (Wang et al., 2021). These studies suggest that mammalian NPs do have a regenerative capability, but it is not sufficient and is not sustainable. Thus, revitalizing and enhancing the regenerative capability of human NP is a legitimate and promising strategy to prevent and treat IDD. This goal may be facilitated by studying the molecular mechanisms that underlie zebrafish NP homeostasis. Supporting this notion, some candidate stem cell or progenitor cell marker genes of mammals are expressed in zebrafish (Table 3), suggesting that the regenerative mechanisms in zebrafish and mammals may have something in common. However, some other mammalian candidate stem cell or progenitor cell marker genes were not expressed in the zebrafish NP. How to correlate the similarities and differences in candidate stem cell gene expression between zebrafish and mammals with their differences in NP maintenance would be an interesting question for future investigation.

### Zebrafish as a model to study the effects of mechanical stresses on the IVD

Mechanical stresses affect IVD homeostasis, and IDD is correlated with excessive physical loads in humans (Iwakiri et al., 2023; Jensen et al., 2012). Although it is not fully understood how mechanical stresses contribute to IDD at the molecular level, it has been shown that compressive and tensile stresses can induce changes in gene expression in NP and AF cells (Hartman et al., 2015; Sowa et al., 2011). The body restriction mold we devised to bend the fish body at various curvatures (Fig. 6D) is similar to the strategy of bending the mouse tail to cause mechanical stresses on the AF and the NP, which resulted in increased cell death and reduced aggrecan expression in the NP matrix (Court et al., 2001). Although not bent as much as the caudal spine, the IVDs in the precaudal vertebrae in zebrafish are also expected to be subjected to compressive and stretching stresses because zebrafish NPs and notochordal strands are connected as one entity that may spread the physical stresses craniocaudally. Because the bending of the spine results in observable conformational changes in the equatorial basal cells and the nearby vacuolated NP cells, the body restriction model is expected to help study the mechanosensation and mechanotransduction in the NP cells in IDD and scoliosis.

### Zebrafish as a genetic and developmental model to study the NP

Genetic factors are involved in IDD, as demonstrated by the obvious predisposition to IDD in chondrodystrophoid but not non-chondrodystrophoid dog breeds (Hansen, 1952) and the differences in IDD development in mice of different genetic backgrounds (Brent et al., 2020). Genetic factors involved in NP degeneration and regeneration can be more conveniently studied in zebrafish thanks to the following four advantages (Choi et al., 2021; Lieschke and Currie, 2007; Teame et al., 2019): First, zebrafish have a lifespan of 2−3 years, and fish at 1.5 ypf normally stop breeding and are already considered aged fish. This lifespan is much shorter than that of many large animal models or humans, thus facilitating aging research. Furthermore, NP fibrosis starts early on in young adults, further shortening the time needed to see degeneration and regeneration effects. Second, zebrafish have high fecundity. They can breed year-round.

Females can lay eggs weekly, and males can mate every two days. A productive female can lay 100−400 eggs each week, making it convenient to perform genetic screens. Third, zebrafish embryos develop externally, making it easy to perform embryonic injections to generate genetically-modified fish lines and screen water-soluble drugs for assessing their therapeutic effects. For example, 30% of embryos injected with CRISPR-based mutagenic reagents can produce gene-specific mutations in germ line cells; 10−30% of transgene-injected fish embryos carry germ line transmission; and 5−10% of embryos injected with CRISPR-based DNA knock-in reagents can produce knock-in fish lines via homologous recombination. Moreover, the external development and the optical transparency of embryos and larvae allow easy observation of NP development. Fourth, it is economical to use zebrafish because they require less housing space. For example, around 135 fish can be kept in a 9-liter fish tank. This drastically reduces maintenance costs. These unique advantages make zebrafish a convenient genetic model to study IVD development, homeostasis, degeneration, and regeneration, thus complementing mammalian models.

## Acknowledgments

XW was supported by the Department of Ophthalmology at the University of Pittsburgh.

